# Synthetic Promoter Design in *Escherichia coli* based on Generative Adversarial Network

**DOI:** 10.1101/563775

**Authors:** Ye Wang, Haochen Wang, Lei Wei, Shuailin Li, Liyang Liu, Xiaowo Wang

**Affiliations:** Ministry of Education Key Laboratory of Bioinformatics; Center for Synthetic and Systems Biology; Bioinformatics Division, Beijing National Research Center for Information Science and Technology; Department of Automation, Tsinghua University, Beijing, 100084, China

**Keywords:** synthetic promoters, generative adversarial network, transcriptional fine-tuning, computational design

## Abstract

Promoter design remains one of the most important considerations in metabolic engineering and synthetic biology applications. Theoretically, there are 4^50^ possible sequences for a 50-nt promoter, of which naturally occurring promoters only make up a small subset. In order to explore the vast potential sequences, we report a novel AI-based framework for *de novo* promoter design in *E. coli.* Guided by the sequence features learned from natural promoter sequences, our framework could explore the potential sequences effectively and design brand new synthetic promoters *in silico.* Specifically, Wasserstein generative adversarial network with gradient penalty (WGAN-GP) and convolutional neural network (CNN) based model were used to design promoters with high activity, and their activities were further tested *in vivo.* As a result, more than 45% of the artificial promoters were demonstrated to be functional and shared no significant sequence similarity with neither natural promoters nor the *E. coli* genome. Our work provides a new approach to design novel functional promoters effectively, indicating the potential for deep learning approaches to be applied into *de novo* genetic element design.

## INTRODUCTION

Well-characterized regulatory elements are indispensable tools for synthetic circuit design and metabolic engineering, which offer enormous potential for industrial biotechnology by producing chemical, medical and material products^*1, 2*^.Promoter is a key regulator in gene expression at transcription level, hence the choice of promoter elements remains an essential consideration in synthetic biology applications^*3*^. Although natural promoters in various organisms are available, they still constrain the transcriptional level to a limited activity range^*4*^. Thus, researchers have proposed several methods to design novel synthetic promoters^*3, 5, 6*^.

Previous studies searching for novel promoters have been mainly focused on mutagenesis^*5, 7, 8*^ and element combination methods^*9-11*^. Methods based on mutagenesis such as constructing random mutation libraries were reported to successfully generate novel synthetic promoters^*12*^. For example, Alper *et al* used error-prone PCR to mutagenize a bacteriophage PL-λ promoter in *E.coli*, resulting in a novel library with 22 functional mutants^*13*^. In addition, element combination strategy such as integrating TF binding sites with promoter backgrounds^*14, 15*^, or combining shorter functional components^*10, 11*^, also have provided some novel promoters for regulating the target genes.

Even though these approaches have certainly generated new promoter elements, they mainly involved the modification of naturally occurring promoters. From the perspective of sequence coding space, the number of possible sequences of a 50-nt prokaryotic promoter is theoretically estimated to be on the order of 4^50^ (∼ 10^30^), not to mention the scale of eukaryotic promoters, which have longer promoter length and more complex structures. Thus, the natural evolutionary process has only sampled an infinitesimal subset from the potential sequences and the vast potential sequence space remains unexplored. Therefore, it is an interesting question whether one could navigate the potential sequence space effectively to find novel functional promoter elements. To achieve a broader range of sequence exploration by a faster and more effective way, better *de novo* sequence design strategy is clear needed.

Recent developments in deep learning methods have provided novel alternative approaches for promoter sequence design. Especially, generative adversarial network (GAN)^*16*^, a deep learning-based generative model, offers a promising way to navigate the sequence space and thus to generate novel samples. GAN model is based on the minimax game between two neural networks, the generator and the discriminator. The discriminator could extract features from real sample groups, and then guides generator to create new samples in the huge potential space^*17*^ GAN has given birth to sets of state-of-art image generation methods^*18-20*^ and showed its ability in generating new images^*21, 22*^. Recently, some variants of GAN have also been used to design probes for protein binding microarrays^*23*^ and synthetic genes coding for antimicrobial peptides^*24*^ as well as drug-like molecular structures^*25-27*^.

Here, we proposed a GAN-based approach for *de nova* promoter sequence design and validated the activities of the generated promoters *in vivo.* Taking natural promoter sequences as inputs, our model automatically extracted the important promoter sequence features. Guided by the extracted features, the generator designed novel artificial sequences automatically from the huge sequence space. These AI-generated promoters mimic key characteristics of the natural promoters such as k-mer frequency, –10 and –35 motifs and their spacing constraint, while show low global sequence similarity to the natural promoters and the *E. coli* genome. More than 45% of the artificial sequences selected by our prediction model could be experimentally validated as functional promoters, and a number of them showed comparable or even higher activities than strong constitutive promoters as well as their strongest mutants.

## RESULTS AND DISCUSSION

### An AI framework for *de novo* promoter design

For the aim of generating functional synthetic promoters, we introduced a deep learning-based framework (Figure 1), including a GAN network for *de nova* promoter generation and a deep convolutional neural network (CNN) for promoter strength prediction. Then activities of the generated artificial promoters were tested by fluorescent protein expression level in *E*. *coli.*

**Figure 1.**
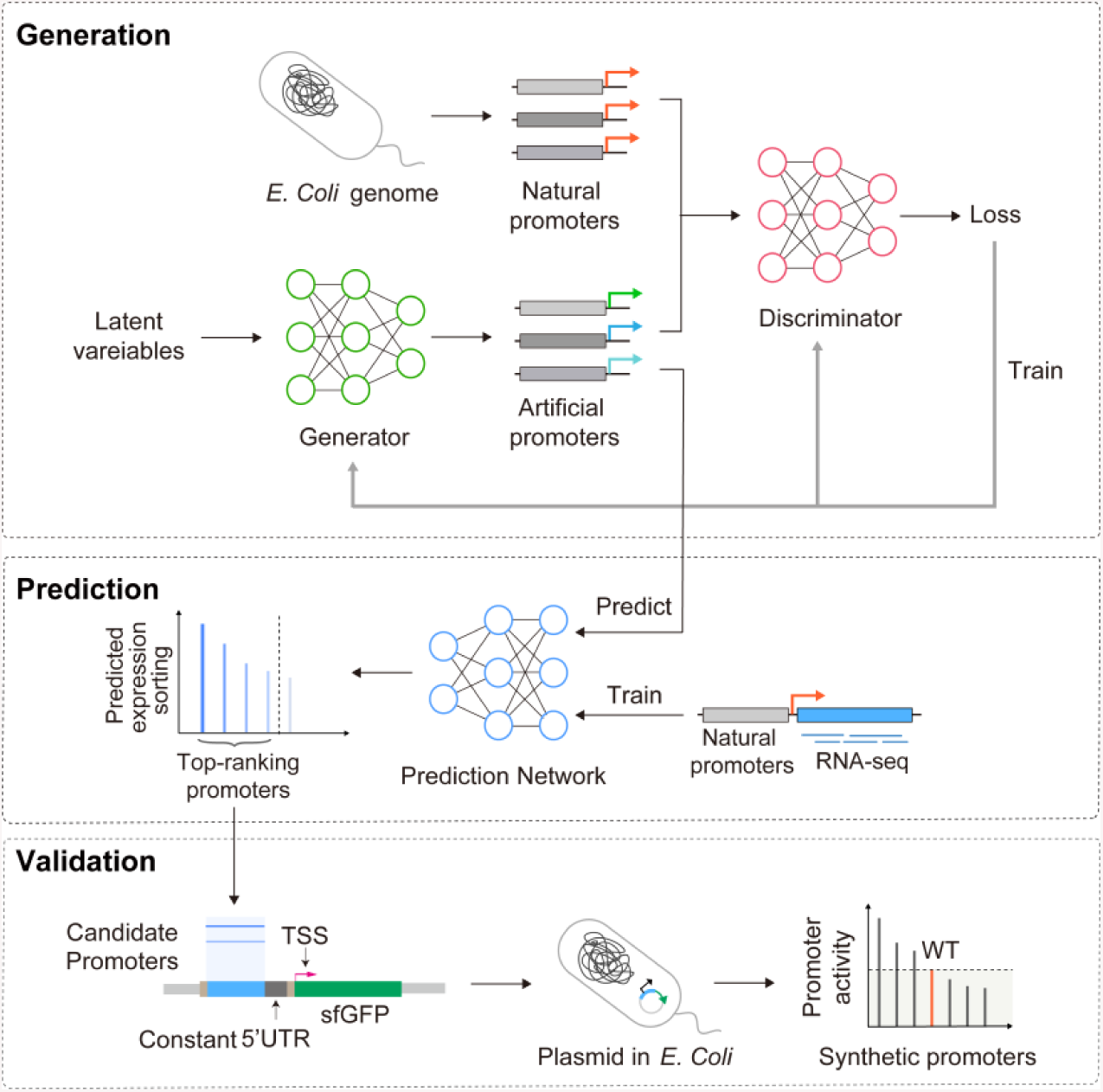
The artificial promoter design approach. In the generation stage, millions of artificial promoters were generated by a GAN model. By activity prediction, candidate promoters with high potential expression level were selected out and were then experimentally validated *in vivo.*

The GAN model has two main adversarial components, the discriminator and generator. The basic workflow of GAN goes as follows: The generator takes samples from the low-dimensional latent variable to generate fake promoters, and the discriminator evaluates the divergence between the current generated fake promoters and the natural promoters. By training the discriminator and generator iteratively, the discriminator gradually learns to act as a more meticulous critic of the current fake promoters. And the generator learns to improve its skills of mimicking the naturally occurring promoters in order to the deceive the discriminator. By producing promoters from the finally trained generator, GAN could help us navigate the potential sequence space to generate new synthetic promoters.

The natural promoters have diverse activities, most of which are lowly expressed in a certain condition. To generate promoters with relatively high activities, a selection module was introduced to predict the gene expression level of the GAN-generated promoters. Specifically, a CNN model was trained by the dataset from Thomason *et al*^*28*^ which contains 14098 promoters in S0bp length with corresponding gene expression levels (Material and Methods). The prediction model structure is shown in Figure S1 Only the promoters predicted with high activities were remained for experimental validation.

Finally, the selected synthetic promoter sequences were used to build the promoter library in *E. coli.* A fixed 5’UTR region was used to control the influence of interaction effects between core transcriptional elements^*29*^. The selected artificial sequences were designed to drive the expression of *sfgfp* gene^*30*^ and their promoter activities were verified *in vivo.*

### The WGAN-GP model for *de novo* promoter generation

At first; we introduced a deep convolutional GAN (DCGAN) based network structure into synthetic promoter design (Figure S2(a)), which was widely used in image generation tasks^*31-33*^. Previous studies on genome sequence pattern learning have shown that convolutional layers could extract motif features and multilayer network could learn the motif combinatorial pattern of specific genomic regions^*34-36*^. The feature learning ability of DCGAN model was analyzed by WebLogo^*37*^ and k-mer frequency. The WebLogo shows independent frequency distribution at each base position and the k-mer frequency represents the high order dependency of nucleotides in promoter sequences. As shown in Figure S2(b), the DCGAN model could partly learn the sequence motif at −10 and −35 region. However, some unexpected base preferences appeared (Figure S2(b)), indicating that the sequences designed by DCGAN model are biased. Besides, the k-mer frequency of DCGAN model promoters differed greatly from the natural promoters (Figure 2(a)) and the k-mer frequency correlation with natural promoters decreased quickly as the k-mer became longer (Figure 2(b)), which illustrated that DCGAN model was not good at capturing longer k-mer frequency features. Such an insufficient feature extraction ability of the DCGAN model would lead to an ineffective exploration of the sequence space, thus the DCGAN model would be inappropriate for promote design.

**Figure 2.**
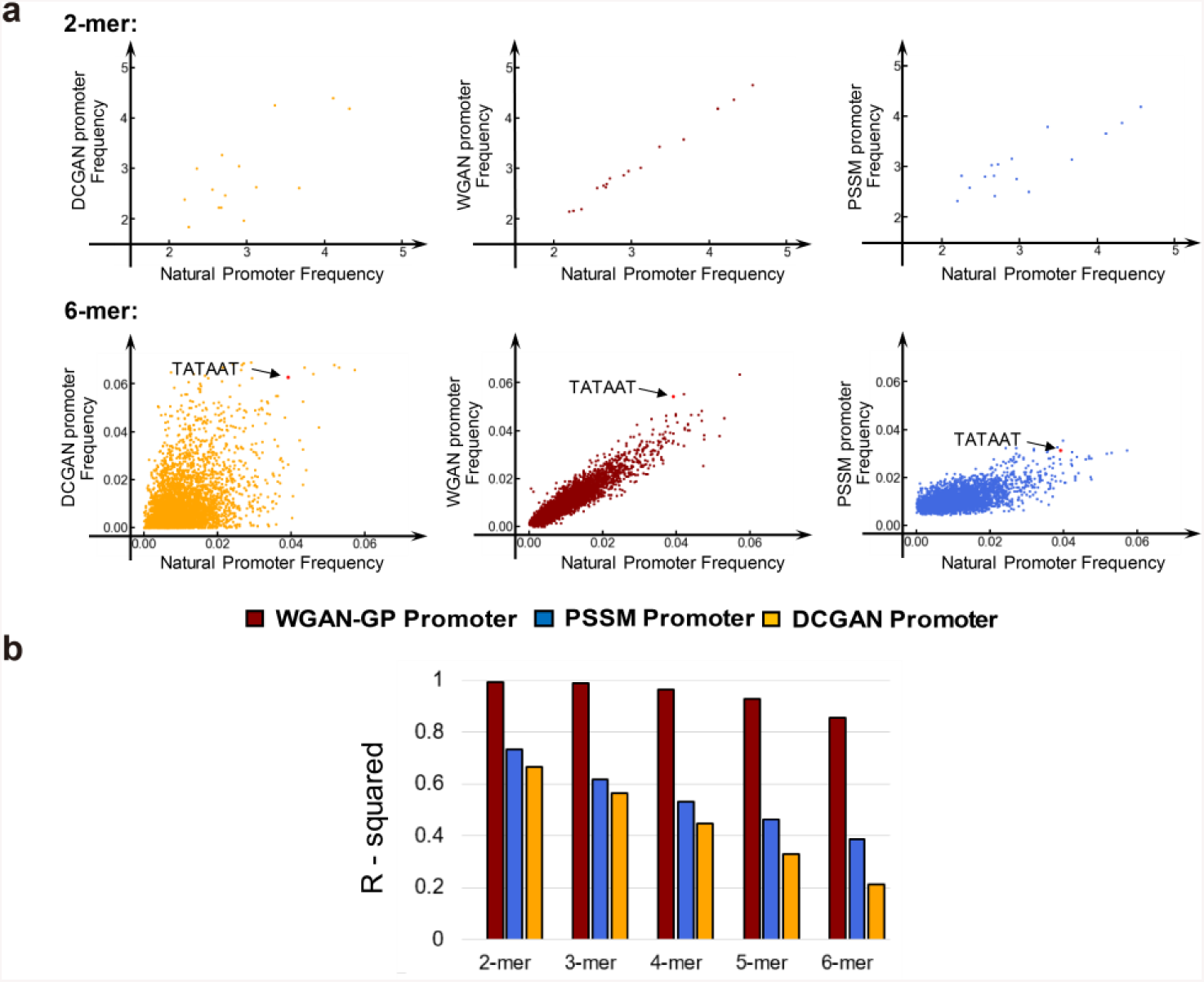
(a) The k-mer scatter plot of natural promoters and generated promoters. Each point represents a certain k-mer. The x and y axes represent the k-mer frequency in natural and generated promoter groups, (b) R-squared value evaluates the correlation of k-mer frequencies (k=2 to 6) between nature promoters and synthetic promoters

Inspired by recent developments of GAN models^*24, 38*^,WGAN-GP language model was then implemented. The WGAN-GP language model was originally designed for nature language generation^*38*^. The network structure is shown in Figure S3. Two main changes were used in WGAN-GP model to solve problems in DCGAN: the introduction of resblock structure^*39*^ and the utilization of earth-moving distance^*40*^.

Resblock is a combination of convolutional layers, with an additional short connection from the resblock start to resblock end. One of the resblock structures are shown in Figure S3. Resblock structure was proposed by He *et al*^*39*^ to handle the gradient degradation problem in deep networks, and enable the deep networks to improve their feature learning abilities. In our work, we found that in synthetic promoter design, resblock structures could decrease the unexpected base preferences compared with the neural network without resblocks. (Figure 3(b)(c)).

**Figure 3.**
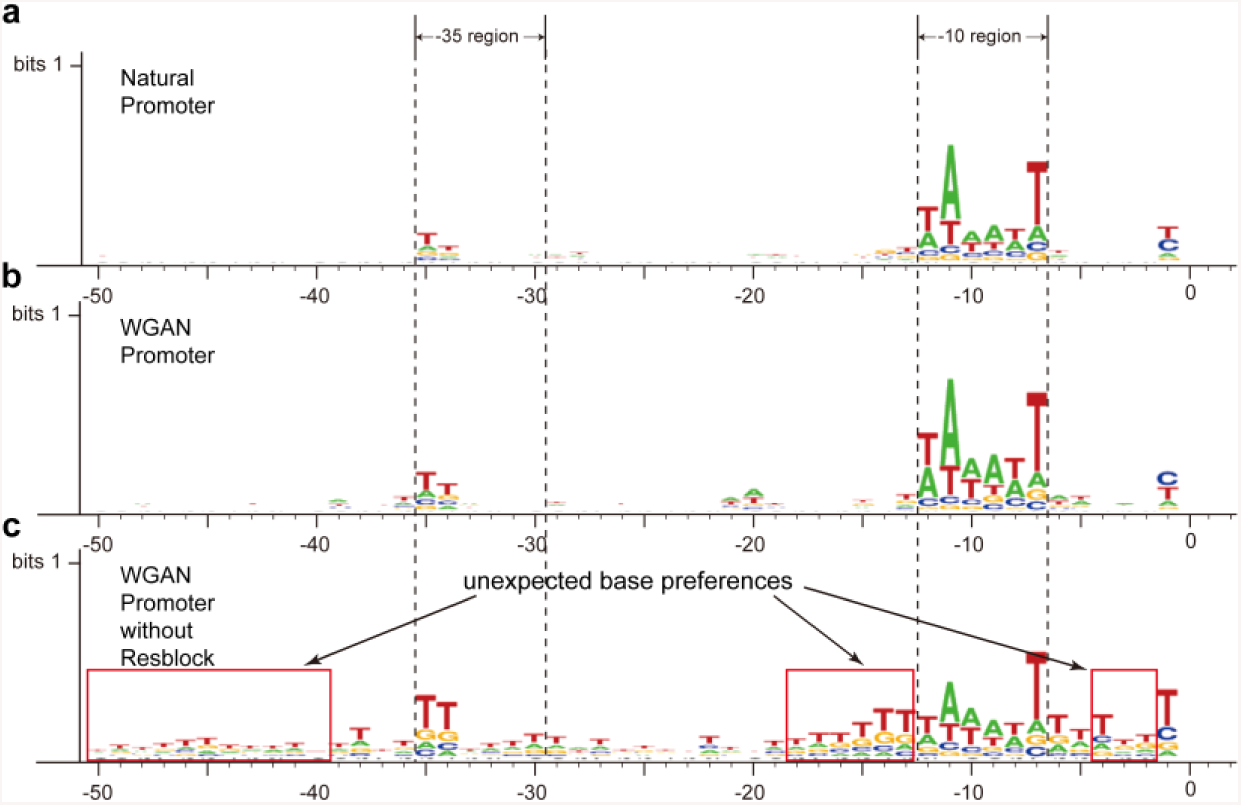
The sequence logo of (a) natural promoters (b) Artificial promoters generated by WGAN-GP with resblock (c) Artificial promoters generated by WGAN-GP without resblock. –10 region and–35 region are annotated on the figures.

Besides, the WGAN-GP model uses the earth-moving distance instead of the <u>Jensen-Shannon</u> divergence^*40*^ to measure the distance between the probability distribution of true samples and generated samples. Unlike the JS divergence, which could not maintain continuous, the earth-moving distance is continuous and provides a usable gradient everywhere^*40*^. Thus, it could progressively train the model until the generated promoter distribution converge to the natural one. By using earth-moving distance, the motif logo of −10 and −35 region was nearly perfectly learned by the WGAN-GP model (Figure 3(a) (b)).

### AI could capture essential promoter sequence features

To explore the characteristics of WGAN-GP generated promoters, the k-mer frequency and the motif spacing constraint were analyzed, and compared with those generated by DCGAN. The position-specific scoring matrix (PSSM) sampling method was also used as a control method to generate bases at each position independently, according to natural promoter base frequencies.

First, we calculated the 2-mer to 6-mer frequency in promoters generated by DCGAN, WGAN-GP and PSSM method (Figure 2(a), Figure S4). Because the synthetic promoters were generated based on the characteristics learnt from natural promoters, we expected that the frequency of crucial k-mers, like Pribnow box (TATAAT) should be well-maintained in the synthetic promoter groups. We found that the occurrence frequency of some common 6-mers, such as TATAAT, decreased in the PSSM method, whereas they are increased in WGAN-GP method (Figure 2(a)). We also compared the top 10 most frequently occurring 6-mers in natural promoters with WGAN-GP, DCGAN and PSSM promoters. As a result, the WGAN shared five 6-mers with natural promoters, with only one in DCGAN promoters as well as PSSM promoters, indicating that the WGAN-GP method had captured important k-mers in natural promoters (Table S1).

To explore the position distribution of the five most frequently occurring 6-mers in natural promoters as well as the well-known Pribnow box (TATAAT), their relative distance to TSS in promoters generated by DCGAN, WGAN-GP and PSSM method were analyzed. As shown in Figure 4(a), the WGAN-GP model outperformed the PSSM method and showed a more similar position distribution with natural promoters, which demonstrated that the WGAN-GP model could learn the k-mer location preference of natural promoters. Besides, the space length between −10 and −35 region was calculated. It was previously reported to be beneficial for RNA polymerase binding if the space length is approximately 16 to 18nts^*41*^. As a result, the space length distribution of WGAN-GP promoter sequences was more centrally distributed in the interval of 16 to 18 region, compared with those of DCGAN and PSSM promoters (Figure 4(c)).

**Figure 4.**
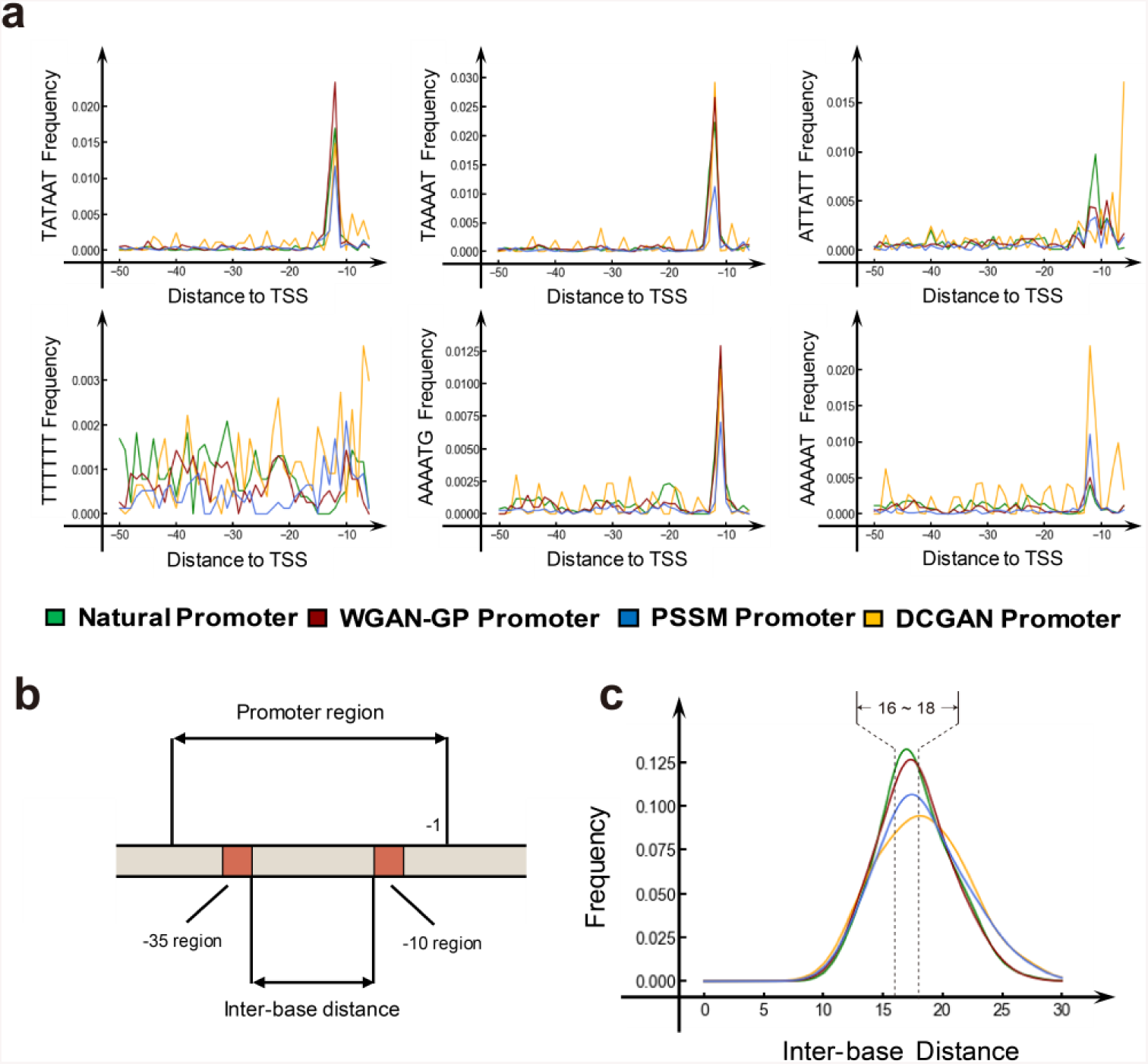
(a) Pribnow box consensus sequence (TATAAT) and top five common 6-mers were selected to illustrate the location learning ability of GAN model. The x axis represents the location relative to TSS, and y axis represents the frequency of 6-mer at this location. (b)The definition of inter based distance between −10 and −35 motifs. (c) The inter base distance distribution between the –10 and –35 motifs. The natural promoters are marked in green, the WGAN-GP, PSSM and DCGAN generated promoters are marked in red, blue, and orange respectively.

The possible explanation for the performance gap between PSSM method and GAN may be that, the PSSM method only focuses on the first-order frequency of bases, whereas the GAN method could learn the high-order dependencies by the powerful feature extraction ability of deep neural networks. In conclusion, the WGAN-GP model showed ability in capturing essential promoter sequence features, including crucial k-mers, k-mer location information and space constraint between −10 and −35 regions.

### The AI-designed promoters show high valid proportion and transcriptional activity

Eighty-three artificial promoters selected by our prediction model were tested in E. *coli.* Besides, two wild-type promoters(BBa_J23119 and Trc1) and their corresponding strong mutants^*42*^ (BBaJ23100, BBa_J23102 and Trc2, Trc3)were also tested. In addition, negative control variants were generated using 5 complete random sequences (Material and Methods). The detailed sequences are provided as Supplementary Table S2.

As a result, 45.8% (38 out of 83) designed promoter sequences showed significantly higher promoter activities than those of random sequences (Benjamini Hochberg, FDR < 5%). Interestingly, three of the synthetic promoters showed comparable or even higher strength than all the six positive control promoters (Figure 5). Most of the functional promoters showed a comparative expression level with the wild-type controls. The promoter activities of all the 83 synthetic promoter were shown in Figure S5.

**Figure 5.**
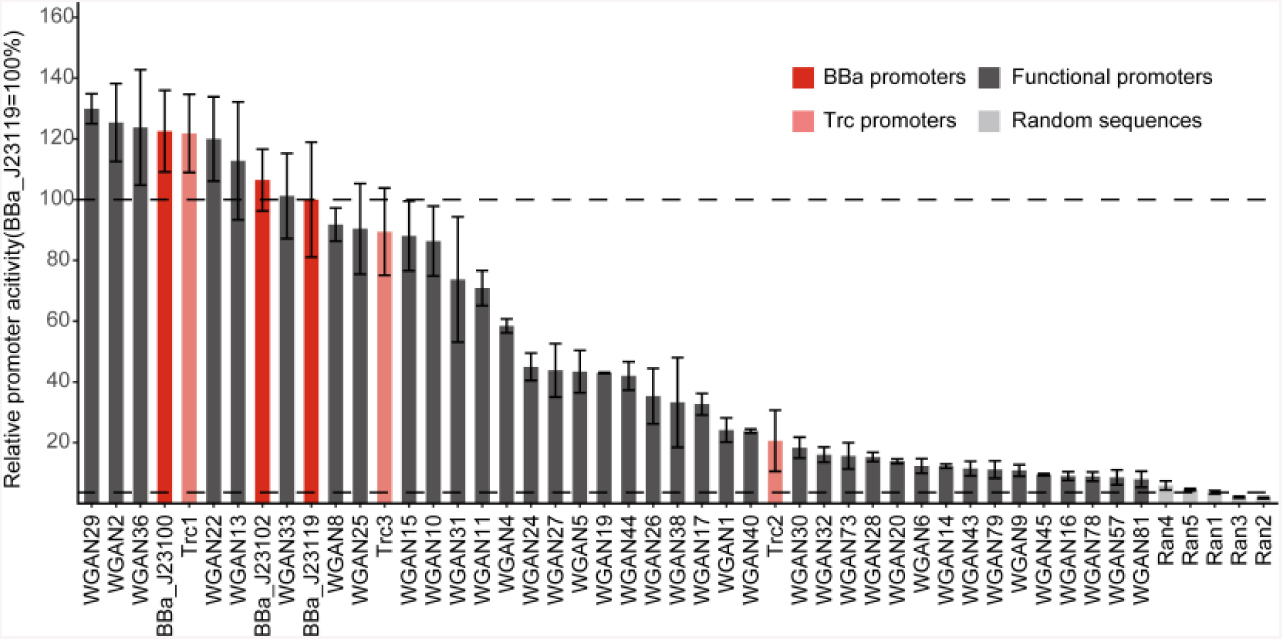
The promoter activity of 38 functional promoters designed by WGAN-GP. BBa_J23119 (dark red) and Trc(light red) are wild-type promoters. BBa_]23100,BBa_]23102 and Trc2,Trc3 are their highly expressed mutants. Rans are five random sequences. The rank of WGAN promoters (dark grey) are the predicted expression level rank obtained from the CNN model. The dashed lines are the 100% baseline represented by BBa_]23119 (high) and average relative activity of five random sequences(low) respectively.

### AI-generated functional elements differ from natural *E. coli* genome sequences

To compare our novel promoters with natural sequences, we analyzed the similarity score between the natural promoters and the AI-generated promoters by CLUSTALW2^*43*^. Specifically, the similarity between our 83 synthetic promoters and the training promoter groups^*28*^ were compared. It turned out that the maximum similarity score is 72%, and the mean is only 62.04% (Figure S6), which means that on average 18 out of total 50 bases in each promoter is different from the natural promoters in training groups.

Besides, a standard nucleotide BLAST searching was also conducted against the whole *E. coli* genome and no significant matches were found. The highest e-value obtained for AI-generated promoters was 0.014, indicating the newly designed promoter sequences are different from the natural genome.

In summary, the essential feature of synthetic promoters, high ratio of functional promoters and the low level of sequence similarity with natural promoters confirmed the effectiveness of WGAN-GP model. These results showed that our framework could design completely new synthetic promoters without copying the original sequences, indicating that the WGAN model could effectively explore the sequence space to find novel promoters in *E. coli.*

### The pros and outlooks of AI-based model in synthetic promoter design

Naturally occurring promoters have evolved for billions of years, but only make up a small subset in the huge potential sequence space. Taking advantage of feature learning ability of GAN models, the functional promoter domain could be learnt in the sequence space. Thus, a great number of synthetic promoters could be designed effectively by navigating the functional promoter domain, which could largely extend functional promoter sequence reservoir.

From the perspective of pattern recognition, the transcriptional machine could be considered as a molecular classifier, which distinguish real promoter sequences from the other genomic regions to initiate transcription. Thus, the synthetic promoters need to have similar properties as natural promoters to recruit the transcriptional machine. Methods based on GAN suit this logic well: the discriminator learns to distinguish the real promoters from the artificial ones, mimicking the role of transcriptional machine. While the generator tries to produce artificial promoters which have similar high order features to natural promoters, resembling the mutation process in nature. However, unlike the naturally occurring mechanisms, in which promoters generally mutant randomly and passively, our generator network could correct its parameters to generate promoters directionally, which speeds up the evolution process.

Specifically, the generator samples from low-dimensional random variables to generate fake promoters. Under the guidance of discriminator, the generator could progressively learn the effective generation strategy to transform random variables into functional promoters. When we input a low-dimensional random variable to the trained generator, it could generate the artificial promoter sequences automatically. This progress provides an effective strategy to explore the vast potential space by learning to natural sequences with important sequence features, avoiding large scale trail-and-error biological experiments.

Our work also indicates the potential application of other deep learning models for element designing tasks. For example, in order to optimize the synthetic elements to specific properties, such as light-responsive promoters^*44*^, other computational methods such as transfer learning^*45*^ and reinforcement learning^*46*^ may be employed. Moreover, other variants of GAN may also be introduced into multi-task element designing^*47*^. For example, the conditional GAN model (*cGAN)*^*48*^, which could generate samples with different properties conditioning on additional information, could be implemented into designing of inducible promoters of distinct inducers, in order to regulate the gene expression in different biological occasions.

Although the selection stage was implemented to predict the target gene expression of promoters, the final successful ratio of functional promoters is still limited. Besides, the relationship between predicted expression level and *in vivo* sfGFP fluorescence is moderate (spearman correlation coefficient= 0.55). The possible explanation may be that: The genome-wide dRNA-seq data in bacteria harbor much noise^41^, which influenced the prediction accuracy of target gene expression level. Thus, if high-throughput fluorescence data paired with promoter sequences in *E. coli* would be available in the future, higher accuracy of prediction model may help increase the valid ratio of functional promoters in our method.

## CONCLUSION

In this study, we conducted *de novo* promoter design based on GAN framework and validated synthetic promoter activities *in vivo.* Our method was benefited from the recent developments in the deep generative model WGAN-GP, which helps to extract high-order base dependence information and generate millions of novel promoters. *In vivo* experimental results suggested that 45.8% of selected novel sequences are functional promoters in *E. coli.* These generated promoters inherent the key features of *E. coli* promoters like the −10 and −35 regions, and have similar k-mer frequency with natural promoters, but also show high sequence difference from the natural promoters and the *E. coli* genome. Such feature could help avoid the genetic instability in the genome context, due to the lower probability of recombination with *E. coli* genome^*9*^.

In summary, we proposed an AI based generative framework for *de novo* design of promoter sequences, which could generate novel promoters that showed high successfully rate by experimentally validation *in vivo.* Our work provided insights into an area of *de novo* synthetic element design, indicating the potential ability for deep learning approaches to explore the sequence space of synthetic elements.

## MATERIALS AND METHODS

### Strains and Cultures

The *E. coli* strain DH5α [F^−^, λ^−^, φ80*lacZ*Δ*M*l5, Δ (*lacZYA*-*argF*), U169, endAl, recAl, *hsdRI7(rk^−^, mk*^*+*^), sup E44, thi^−^, *gyrA96*, relAl, phoA] was used as host organism and cultivated in Luria-Bertani (LB) media supplemented with 100 µg/mL kanamycin at 37 °C for promoter activity validation.

### Promoter library plasmid construction

All promoter constructs were carried on the high-copy number vector Tl.3, driving the expression of sfGFP from the reporter gene *sfgfp*. Promoter and 5’UTR sequences are displayed in Table S2. The forward primer of 5’UTR fragments (B0030) contained an EcoRI site and reverse primers contained an XbaI site. The putative Shine-Dalgarno sequence in 5’UTR was placed 7nt upward of the original TSS region. Annealing reactions were performed by incubating the complementary oligonucleotides at 95 °C for 3 min (2 µl of 100 µM forward and reverse oligonucleotides in sterile water) followed by 95 °C for 1 min each cycle with 57 cycles and cooling to 4°C for storage. The phosphorylation of annealed oligonucleotides was performed using T4 polynucleotide kinase (PNK from NEB) with ATP for 1 hour. Then the 5’UTR oligonucleotides were cloned into the EcoRI-XbaI sites in Tl.3 constructs by T4 ligase catalyzing reaction. The recombinant plasmids with 5’UTR sequences were verified by sequencing. The adenine instead of thymine were designed in 5’UTR oligonucleotides at XbaI site, therefore the original XbaI site could not be digested in following steps. The Tl.3 constructs with 5’UTR oligonucleotides were digested with restriction enzymes EcoRI and XbaI, and designed promoter oligonucleotides flanked with the same MCS were cloned into the new EcoRI-XbaI sites. Six positive controls promoters, five random baseline promoters (with GC content near 50%) and two blank control plasmids were also tested for promoter activity. All of the reporter plasmids were verified by sequencing.

### In vivo promoter assays

Assay strains were stored as glycerol stocks in sterile centrifuge tubes (1.5 ml). Cultures were grown in plates containing 5ml LB medium with kanamycin, inoculating from glycerol stocks using a sterilized metal pinner. Monoclonal selections were grown overnight (16 h) in 96-well U-bottom deep-well plate covered with sterile breathable sealing film (sterile sealing films; Axygen) at 30°C with shaking at 300 rpm on orbital shaker. The following day the overnight cultures were diluted 1:100 into a final volume of 1.5ml of fresh medium with appropriate kanamycin and grown for another 8 hours. In total 200ul cultures were added in clear bottom black plates, and repeated measurements of the optical density at 600 nm(OD600) and fluorescence (relative fluorescence units [RFU]; excitation at 485 nm and emission at 510 nm) were performed by microplate reader-incubator-shaker (Thermo). All experiments were repeated at least three times, and changing the well positions of strains were also conducted in the microplates during three repeated experiments to avoid any local position effects.

### Establishing a Baseline Expression Level of Functional Promoters

Five control variants were generated in which the synthetic promoters were replaced by complete random sequences, with the GC content was controlled at near 50%. The resulting expression levels measured the basal expression of the *sfgfp* gene given that there is enough sequence space upstream of the protein coding sequence for transcription initiation. This experiment was conducted for testing the basic transcription baseline in our system. The average relative promoter activity of the three control variants was 3.6% of the average of the wild type constitutive promoter BBa_JS23119 (BioBrick part in iGEM Registry of Standard Biological Parts). Synthetic promoters with significant higher relative promoter activity were marked as functional promoters, which was selected by Student’s t-test with baseline sequences (Benjamini Hochberg, FDR < 5%).

Six positive control promoter sequences were also used in the present work, including two different types of wild-type promoters: BBa_J23119 and Ptrc, with two of their corresponding mutants (BBa_J23100, BBa_J23102 andPtrc_m0l0,Ptrc_m004), which have shown highest expression in previously reported mutation library^*42*^. BBa_J23100, BBa_J23102 were obtained from the BioBrick part in iGEM Registry of Standard Biological Parts and Ptrc, Ptrc_m010 and Ptrc_m004 were obtained from previous research^*42*^.

Two repeated blank control variants were designed by replacing the synthetic promoter sequences with a 10nt random sequence (GGGCCTGTA), which could not provide enough length for RNA polymerase to bind on the upward sequence of protein coding region. We performed the blank control reporter to avoid the background noise in the system and this value was also used for relative promoter activity calculating.

### Promoter strength analysis

Culture background fluorescence was determined using blank control variant plasmid carrying the same 5’UTR sequence without promoter sequence. This control vector strain was grown and assayed under the same condition with the promoter library. The fluorescence of tested promoters subtracted background fluorescence from the blank control strain under for each reporter vector was calculated as^*42*^

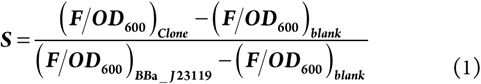

The bacteria density was measured by average optical density at 600 nm (OD600) versus time(h^−1^). Final reported promoter activities were calculated by taking the average of three independent biological experiments.

### The training promoter samples

The training dataset contains in total 14098 experimentally identified promoters in the *E. coli* K12 MG1655 genome. As the description of the dataset, most of the promoters recognized in this dataset were σ^*70*^ promoters.

The promoter sequence is defined as 50bp long upstream of transcription start site (TSS), which could include key motifs such as −10 and −35 region, but not too long to exclude unnecessary sequences^*49*^.

### The brief introduction of GAN

Generative adversarial network (GAN) has achieved impressive results in the fields of nature image generation^*32, 50-52*^,image-to-image translation ^*19, 31, 53*^, super resolution image creation^*18*^. Recent studies show that by learning the high dimensional representation of real sample group, GAN model has the ability to generating millions of completely new samples. More specifically, GAN model contains two parts, the generator and the discriminator. The aim of generator is to generate fake samples that could not be distinguished with real samples by the discriminator, whereas the aim of discriminator is learning to classify the fake samples and the real samples. With iteratively optimizing two counter objectives, the generator could progressively generate fake samples just as real samples.

Mathematically, in the original version of *GAN*^*16*^ the aim of discriminator is as follows:

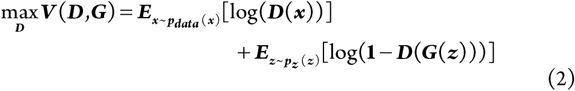

Here, ***D*(*x)*** represents the classification result of real samples. ***z*** represents input of generator, which usually is the low dimension variable. ***D*(*z*)** represents generated fake samples, and ***D*(*G*(*z*))** represent the classification result of fake samples. The first part demonstrates the discriminator should recognize real samples better, and the second part means it should recognize generated samples better. On the contrary, the aim of generator is:

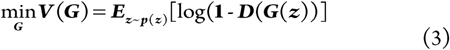

It means that the generator aims to generate images that has the high possibility to fool the discriminator. Thus, GAN model is playing a minmax game by the generator and discriminator, which could be shown in the value function:

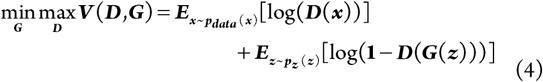

### WGAN-GP language model

The eq.(4) implies that the model is minimizing the JS divergence between the real and fake samples distribution when the discriminator is optimal^*16*^. However, when two distribution have supports are disjoint or lie on low dimension manifolds, such JS divergence could not iteratively make the model generate the correct samples^*54*^. Thus, M Arjovsky *et al* used earth moving distance instead of JS divergence ^*40*^ to measure the distance between two distributions, and such GAN model is called WGAN. Then, Gulrajani I *et al* added a gradient penalty into WGAN value function to solve the optimization difficulties in the WGAN^*38*^. The WGAN-GP models have recovered the instability of original GAN model and achieved great results in the image and nature language generation. In our work, we used WGAN-GP language model in our promoter generation task. Such model has been used in the DNA sequence generation^*24*^, which verifies the effectiveness of the model towards automatic design of promoters.

### Find the −10 and −35 motif by PSSM matrix

We first counted the occurrence possibility of each position of promoter sequences to calculate PSSM matrix, then used logit function,i.e. log ***P*_*ij*_ */b*_*i*_**,where ***P***_*ij*_ implies the element in the PSSM matrix and ***b*_*i*_** implies the background distribution. Here, we selected background by calculating the occurrence possibility of T,C,G,A in the whole dataset. The threshold that recognized the motif was set to 3, meanwhile the — 10 and —35 motif finding region was restricted to −1 ∼ −17 and −40∼ −25.

### Top 6-mer position distribution relative to TSS

The position distribution of top 20 most frequently occurring 6-mers in natural promoters were shown in Supplementary Figure S7.

#### Several important parts in the GAN network

1. Convolution layers: The convolution layer is the core part in the promoter feature learning. It contains a great deal of convolution kernels, and each kernel could learn a certain kind of motif features. The parameters of each kernel could be learned during the iteratively training of GAN model. With the combination of several Convolution layers, the whole neuron network could get the information of the promoter sequences in different sizes. The utilization of convolution layer could guide the generator to produce the certain motifs in the generated promoters, and guide the discriminator to better classify the samples.

2. Deconvolution layers: The input of the generator is a low dimension vector whereas the output is a relatively higher dimensional vector. Thus, we need deconvolution layers in DCGAN network to accomplish the mapping from the lower dimensional input to the higher dimensional output.

3. Full connection layer: This kind of layer is the most common layer in the neuron networks, it could achieve linear mapping from one vector to another vector. The convolution layer could be treated as the optimization version of full connection layer, and we use convolution layer more common than the full connection layer.

4. Activation function: The activation function could dramatically influence the generated results. The bad choice of activation function may induce to mode collapse problem. In our DCGAN model, we introduced tanh as the activation function of the intermediate layer.

And we used the softmax as the activation function of output layer of generator. We used softmax rather than tanh as activation function because the appearance of A means the absence of T, C and G, which means the four bases are mutually exclusive. For the output layer of discriminator, we used sigmoid as the activation function in DCGAN model.

### Encoding mode of promoter sequence

Promoter sequences with length N was represented as matrix, where the length N equaled to 50. T, A, C, G was encoded as [1,0,0,0], [0,1,0,0], [0,0,1,0], and [0,0,0,1] respectively.

### Model training of GAN models and CNN prediction model

We used all the promoter sequences in the dataset described above as the real samples. In the DCGAN model, the input of the generator was the uniform distribution random variable. The batch size was set to 128, the iteration times was set to 100 and we used stochastic gradient descent as the optimization method of our model.

Different from the DCGAN, in the WGAN-GP model, we sampled from standard normal distribution as the input variable of generator. The batch size was set to 32 and we trained our network for 160 epochs. We found that the best result is around 12 epochs, so we selected synthetic promoters from that range of iterations. Notice that for giving the best play of the WGAN-GP model, we train 5 times discriminator and one time generator in each batch training. The optimizer used Adam with learning rate equaled to 0.0001, beta1 equaled to 0.5 and beta2 equaled to 0.9.

In the prediction model, the training dataset was from the Thomason *et al* which contains 14098 promoters with corresponding gene expression level. The batch size was set to 128, and we used stochastic gradient descent as the optimization method. We trained this model with 9000 samples as training set, 1000 samples as validation set and others as testing set. Notice that we used kernel size equaled to seven to capture the motif like −10, –35 motif in the promoter region.

## ASSOCIATED CONTENT

### Supporting Information

This material is available free of charge via the Internet at http://pubs.acs.org:

The supporting information (PDF) includes Table S1-S2 and Figures S1-S7: Table S1: List of top 10 most frequently occurring 6-mers in natural promoters; Table S2: Promoters and S’UTR sequences used in present work; Figure S1: Network structure of CNN based predictor; Figure S2: Network structure and sequence logo of DCGAN; Figure S3: Network structure of WGAN-GP; Figure S4: The correlation scatter plot of k-mer frequency of natural promoters and DCGAN, WGAN-GP, PSSM promoters. Figure SS: Promoter activity of83 synthetic promoters design *in silica.* Figure S6: The similarity score distribution of random sequences and DCGAN, WGAN-GP, PSSM promoters. Figure S7: The position distribution of top 20 most frequently occurring 6-mers. Plasmid backbone map in GenBank format.

## ABBREVIATIONS

GAN: Generative adversarial network
DCGAN: Deep Convolutional GAN
WGAN-GP: Wasserstein GAN with gradient penalty

## REFERENCES

(1) Lynch, S. A., and Gill, R. T. (2012) Synthetic biology: new strategies for directing design, Metabolic engineering 14, 205–211.

(2) Arundel, A., and Sawaya, D. (2009) The bioeconomy to 2030: Designing a policy agenda.

(3) Gilman, J., and Love, J. (2016) Synthetic promoter design for new microbial chassis, Biochemical Society transactions 44, 731–737.

(4) Blazeck, J., and Alper, H. S. (2013) Promoter engineering: recent advances in controlling transcription at the most fundamental level, Biotechnology journal 8, 46–58.

(5) Guiziou, S., Sauveplane, V., Chang, H. J., Clerte, C., Declerck, N., Jules, M., and Bonnet, J. (2016) A part toolbox to tune genetic expression in Bacillus subtilis, Nucleic acids research 44, 7495–7508.

(6) De Mey, M., Maertens, J., Lequeux, G. J., Soetaert, W. K., and Vandamme, E. J. (2007) Construction and model-based analysis of a promoter library for E. coli: an indispensable tool for metabolic engineering, BMC biotechnology 7, 34.

(7) Nevoigt, E., Kohnke, J., Fischer, C. R., Alper, H., Stahl, U., and Stephanopoulos, G. (2006) Engineering of promoter replacement cassettes for fine-tuning of gene expression in Saccharomyces cerevisiae, Appl Environ Microbiol 72, 5266–5273.

(8) Du, J., Yuan, Y., Si, T., Lian, J., and Zhao, H. (2012) Customized optimization of metabolic pathways by combinatorial transcriptional engineering, Nucleic acids research 40, e142.

(9) Portela, R. M., Vogl, T., Kniely, C., Fischer, J. E., Oliveira, R., and Glieder, A. (2017) Synthetic Core Promoters as Universal Parts for Fine-Tuning Expression in Different Yeast Species, ACS synthetic biology 6, 471–484.

(10) Blazeck, J., Liu, L., Redden, H., and Alper, H. (2011) Tuning gene expression in Yarrowia lipolytica by a hybrid promoter approach, Appl Environ Microbiol 77, 7905–7914.

(11) Blazeck, J., Garg, R., Reed, B., and Alper, H. S. (2012) Controlling promoter strength and regulation in Saccharomyces cerevisiae using synthetic hybrid promoters, Biotechnology and bioengineering 109, 2884–2895.

(12) Yim, S. S., An, S. J., Kang, M., Lee, J., and Jeong, K. J. (2013) Isolation of fully synthetic promoters for high-level gene expression in Corynebacterium glutamicum, Biotechnology and bioengineering 110, 2959–2969.

(13) Alper, H., Fischer, C., Nevoigt, E., and Stephanopoulos, G. (2005) Tuning genetic control through promoter engineering, Proceedings of the National Academy of Sciences 102, 12678–12683.

(14) Vogl, T., Ruth, C., Pitzer, J., Kickenweiz, T., and Glieder, A. (2014) Synthetic core promoters for Pichia pastoris, ACS synthetic biology 3, 188–191.

(15) Weingarten-Gabbay, S., Nir, R., Lubliner, S., Sharon, E., Kalma, Y., Weinberger, A., and Segal, E. (2019) Systematic interrogation of human promoters, Genome research 29, 171–183.

(16) Goodfellow, I., Pouget-Abadie, J., Mirza, M., Xu, B., Warde-Farley, D., Ozair, S., Courville, A., and Bengio, Y. (2014) Generative adversarial nets, In Advances in neural information processing systems, pp 2672–2680.

(17) Gama-Castro, S., Salgado, H., Santos-Zavaleta, A., Ledezma-Tejeida, D., Muniz-Rascado, L., Garcia-Sotelo, J. S., Alquicira-Hernandez, K., Martinez-Flores, I., Pannier, L., Castro-Mondragon, J. A., Medina-Rivera, A., Solano-Lira, H., Bonavides-Martinez, C., Perez-Rueda, E., Alquicira-Hernandez, S., Porron-Sotelo, L., Lopez-Fuentes, A., Hernandez-Koutoucheva, A., Del Moral-Chavez, V., Rinaldi, F., and Collado-Vides, J. (2016) RegulonDB version 9.0: high-level integration of gene regulation, coexpression, motif clustering and beyond, Nucleic acids research 44, D133–143.

(18) Ledig, C., Theis, L., Huszar, F., Caballero, J., Cunningham, A., Acosta, A., Aitken, A. P., Tejani, A., Totz, J., and Wang, Z. (2017) Photo-Realistic Single Image Super-Resolution Using a Generative Adversarial Network, computer vision and pattern recognition, 105–114.

(19) Zhu, J.-Y., Park, T., Isola, P., and Efros, A. A. (2017) Unpaired image-to-image translation using cycle-consistent adversarial networks, In Proceedings of the IEEE International Conference on Computer Vision, pp 2223–2232.

(20) Li, C., and Wand, M. (2016) Precomputed real-time texture synthesis with markovian generative adversarial networks, In European Conference on Computer Vision, pp 702–716, Springer.

(21) Denton, E. L., Chintala, S., and Fergus, R. (2015) Deep generative image models using a, In Advances in neural information processing systems, pp 1486–1494.

(22) Yang, J., Kannan, A., Batra, D., and Parikh, D. (2017) Lr-gan: Layered recursive generative adversarial networks for image generation, arXiv preprint arXiv:1703.01560.

(23) Gupta, A., and Zou, J. (2019) Feedback GAN for DNA optimizes protein functions, Nature Machine Intelligence 1, 105.

(24) Killoran, N., Lee, L. J., Delong, A., Duvenaud, D., and Frey, B. J. (2017) Generating and designing DNA with deep generative models, arXiv preprint arXiv:1712.06148.

(25) Putin, E., Asadulaev, A., Ivanenkov, Y., Aladinskiy, V., Sanchez-Lengeling, B., Aspuru-Guzik, A., and Zhavoronkov, A. (2018) Reinforced adversarial neural computer for de novo molecular design, Journal of chemical information and modeling 58, 1194–1204.

(26) Kadurin, A., Nikolenko, S., Khrabrov, K., Aliper, A., and Zhavoronkov, A. (2017) druGAN: an advanced generative adversarial autoencoder model for de novo generation of new molecules with desired molecular properties in silico, Molecular pharmaceutics 14, 3098–3104.

(27) De Cao, N., and Kipf, T. (2018) MolGAN: An implicit generative model for small molecular graphs, arXiv preprint arXiv:1805.11973.

(28) Thomason, M. K., Bischler, T., Eisenbart, S. K., Forstner, K. U., Zhang, A., Herbig, A., Nieselt, K., Sharma, C. M., and Storz, G. (2015) Global transcriptional start site mapping using differential RNA sequencing reveals novel antisense RNAs in Escherichia coli, Journal of bacteriology 197, 18–28.

(29) Davis, J. H., Rubin, A. J., and Sauer, R. T. (2011) Design, construction and characterization of a set of insulated bacterial promoters, Nucleic acids research 39, 1131–1141.

(30) Pédelacq, J.-D., Cabantous, S., Tran, T., Terwilliger, T. C., and Waldo, G. S. (2006) Engineering and characterization of a superfolder green fluorescent protein, Nature biotechnology 24, 79.

(31) Isola, P., Zhu, J.-Y., Zhou, T., and Efros, A. A. (2017) Image-to-image translation with conditional adversarial networks, arXiv preprint.

(32) Radford, A., Metz, L., and Chintala, S. (2015) Unsupervised representation learning with deep convolutional generative adversarial networks, arXiv preprint arXiv:1511.06434.

(33) Yoo, D., Kim, N., Park, S., Paek, A. S., and Kweon, I. S. (2016) Pixel-level domain transfer, In European Conference on Computer Vision, pp 517–532, Springer.

(34) Quang, D., and Xie, X. (2016) DanQ: a hybrid convolutional and recurrent deep neural network for quantifying the function of DNA sequences, Nucleic acids research 44, e107–e107.

(35) Zeng, H., Edwards, M. D., Liu, G., and Gifford, D. K. (2016) Convolutional neural network architectures for predicting DNA–protein binding, Bioinformatics 32, i121–i127.

(36) Kelley, D. R., Snoek, J., and Rinn, J. L. (2016) Basset: learning the regulatory code of the accessible genome with deep convolutional neural networks, Genome research.

(37) Crooks, G. E., Hon, G., Chandonia, J.-M., and Brenner, S. E. (2004) WebLogo: a sequence logo generator, Genome research 14, 1188–1190.

(38) Gulrajani, I., Ahmed, F., Arjovsky, M., Dumoulin, V., and Courville, A. C. (2017) Improved training of wasserstein gans, In Advances in Neural Information Processing Systems, pp 5767–5777.

(39) He, K., Zhang, X., Ren, S., and Sun, J. (2016) Deep residual learning for image recognition, In Proceedings of the IEEE conference on computer vision and pattern recognition, pp 770–778.

(40) Arjovsky, M., Chintala, S., and Bottou, L. (2017) Wasserstein gan, arXiv preprint arXiv:1701.07875.

(41) Harley, C. B., and Reynolds, R. P. (1987) Analysis of E. coli pormoter sequences, Nucleic acids research 15, 2343–2361.

(42) Semsey, S., Meng, H., Wang, J., Xiong, Z., Xu, F., Zhao, G., and Wang, Y. (2013) Quantitative Design of Regulatory Elements Based on High-Precision Strength Prediction Using Artificial Neural Network, PLoS ONE 8, e60288.

(43) Larkin, M. A., Blackshields, G., Brown, N., Chenna, R., McGettigan, P. A., McWilliam, H., Valentin, F., Wallace, I. M., Wilm, A., and Lopez, R. (2007) Clustal W and Clustal X version 2.0, Bioinformatics 23, 2947–2948.

(44) Horner, M., Muller, K., and Weber, W. (2017) Light-Responsive Promoters, Methods in molecular biology (Clifton, N.J.) 1651, 173–186.

(45) Segler, M. H., Kogej, T., Tyrchan, C., and Waller, M. P. (2017) Generating focused molecule libraries for drug discovery with recurrent neural networks, ACS central science 4, 120–131.

(46) Popova, M., Isayev, O., and Tropsha, A. (2018) Deep reinforcement learning for de novo drug design, Science advances 4, eaap7885.

(47) Li, Y., Zhang, L., and Liu, Z. (2018) Multi-objective de novo drug design with conditional graph generative model, Journal of cheminformatics 10, 33.

(48) Mirza, M., and Osindero, S. (2014) Conditional generative adversarial nets, arXiv preprint arXiv:1411.1784.

(49) Kim, D., Hong, J. S.-J., Qiu, Y., Nagarajan, H., Seo, J.-H., Cho, B.-K., Tsai, S.-F., and Palsson, B. Ø. (2012) Comparative analysis of regulatory elements between Escherichia coli and Klebsiella pneumoniae by genome-wide transcription start site profiling, PLoS genetics 8, e1002867.

(50) Brock, A., Donahue, J., and Simonyan, K. (2018) Large scale gan training for high fidelity natural image synthesis, arXiv preprint arXiv:1809.11096

(51) Zhang, H., Goodfellow, I., Metaxas, D., and Odena, A. (2018) Self-attention generative adversarial networks, arXiv preprint arXiv:1805.08318.

(52) Miyato, T., Kataoka, T., Koyama, M., and Yoshida, Y. (2018) Spectral normalization for generative adversarial networks, arXiv preprint arXiv:1802.05957.

(53) Choi, Y., Choi, M., Kim, M., Ha, J.-W., Kim, S., and Choo, J. (2018) Stargan: Unified generative adversarial networks for multi-domain image-to-image translation, In Proceedings of the IEEE Conference on Computer Vision and Pattern Recognition, pp 8789–8797.

(54) Arjovsky, M., and Bottou, L. (2017) Towards principled methods for training generative adversarial networks, arXiv preprint arXiv:1701.04862.

