## Supplemental Information for "Synthetic Promoter Design in *Escherichia coli* based on Generative Adversarial Network"

Table S1. The top 10 most frequently occurring 6-mers in natural, WGAN, PSSM and DCGAN promoters are shown in the table. The 6-mers shared with natural promoters are marked in green.

| Natural Promoter | WGAN promoter | PSSM promoter | DCGAN promoter |
|---|---|---|---|
| TAAAAT | TAAAAT | TTAATA | AAAATT |
| ATTATT | TTTATT | TTAAAA | AATATT |
| TTTTTT | TATAAT | AAAAAT | TAAATT |
| AAAATG | TATTAT | TAATAA | AAATTT |
| AAAAAT | ATAAAA | TAAAAA | TATATT |
| AAAAAA | ATTTTT | ATAATT | TTTAAT |
| AATAAT | ATAATG | TTAAGA | AATTTT |
| TATTAT | AAAATT | AAAATT | TAAAAT |
| ATTTTT | TTATTT | TTTATT | TATAAT |
| ATAATG | ATTATT | TTATTA | TAATAT |

Table S2. List of promoters and 5'UTR primer sequences used in constructs of present work with corresponding lengths. The restriction sites (EcoRI and xbaI) are highlighted in purple, and the putative Shine-Dalgarno sequence in 5'UTR sequence is highlighted in blue.

| ID | 5'UTR | Sequence | Length |
|---|---|---|---|
| **U1** | B0030 | AATTCGGGCTCTGTATCTAGAATTAAAGAGGAGAAAA | 15+22 |
| **ID** | **Promoters** | **Sequence** | **Length** |
| **Pc1** | BBa_J23119 | AATTCTTGACAGCTAGCTCAGTCCTAGGTATAATGCTAGCT | 35+6 |
| **Pc2** | BBa_J23100 | AATTCTTGACGGCTAGCTCAGTCCTAGGTACAGTGCTAGCT | 35+6 |
| **Pc3** | BBa_J23102 | AATTCTTGACAGCTAGCTCAGTCCTAGGTACTGTGCTAGCT | 35+6 |
| **Pc4** | Ptrc | AATTCTTGACAATTAATCATCCGGCTCGTATAATGTGTGGAATTGTGAGT | 44+6 |
| **Pc5** | Ptrc_m010 | AATTCTTGACAGTTAATCATCCGGCCCGTACAGTGTGTGGGACTGTGAGT | 44+6 |
| **Pc6** | Ptrc_m004 | AATTCTTGACAATTGGTCATCCGGCTCGTATAATGTGTGGAACTGTGAGT | 44+6 |
| **Ran1** | | AATTCTCTAACATTATATACCCCGCTGAATAAAGGCAGGCCACACTGTTGCGTCGT | 50+6 |
| **Ran2** | | AATTCATTTCAATGCCGAATCGATCCTGTCCTGTTCACCCTCAACCGATAGGGCTT | 50+6 |
| **Ran3** | | AATTCAATATAGTGGAGTGAGCCCTTTCCTCTTCTAGTCAAGTGTCGAGCGGACAT | 50+6 |
| **Ran4** | | AATTCCCGGGGATATACTAGATATTGCGCTGTTGTCTTTCGGAGAAACGGTGGCCT | 50+6 |
| **Ran5** | | AATTCACTGAACTCGTTAAGACTTTGCATGTCACGGAAGAGAGGTGGCAATCCGCT | 50+6 |
| **WGAN1** | | AATTCAGTCAACATTAAAAGTTAAAATAGTTGCTATGATGTGCTAAGGTAAATAAT | 50+6 |
| **WGAN2** | | AATTCGGTCAACAATTATAGTTGATTATCGCGTAAAAGTATGCTACCCTTAAGTTT | 50+6 |
| **WGAN3** | | AATTCTTCATAATTTCGACTTGTCATATTTGCGTAAACTTACGATTTAACCGAGCT | 50+6 |
| **WGAN4** | | AATTCATTCATGCTGATGGATTTGAAATATAGCGATTATTTGTTATGCTAGGCACT | 50+6 |
| **WGAN5** | | AATTCCCAGCCACTTCCCCTTTAAAAAAATCGGGTATACGCGCTATACTATGAATT | 50+6 |
| **WGAN6** | | AATTCTGTTTAACCTTTGGTGTTAACTGTCAGATCTTGTGCGCTATACTTAAATTT | 50+6 |
| **WGAN7** | | AATTCAAAGAAAAAGCTTGATTAAAGTTGAGAAAAATAGGGCCTATGATTCGGCTT | 50+6 |
| **WGAN8** | | AATTCGTCCTGTTAAATCCTTTAAAGATAGAGCGTTTCGTGCTATGCTACGCATCT | 50+6 |
| **WGAN9** | | AATTCTGTTTAAATTTTGATTTAAGTCTTATCGGAGGCTATGCTATTTGCGATTTT | 50+6 |
| **WGAN10** | | AATTCGAAATCGTGTAAACGTTGGATTATAGTTGCATTTATGCTATACTGTTACCT | 50+6 |

| | | | |
|---|---|---|---|
| **WGAN11** | | AATTCCCTTTAGAGAAATACGTTGAATTTGAGGCGTGAGTGACTATACTAAAGGGT | 50+6 |
| **WGAN12** | | AATTCTTTTTAAAAACTTACTTTAAGCTTGAGCGAAAAGGCATTATTAAAGAGACT | 50+6 |
| **WGAN13** | | AATTCTTTTTGACGCATAAGTTGTGAAATCGTGTAATTTCTGCTATGCTAGGTTCT | 50+6 |
| **WGAN14** | | AATTCAAAAAGTGTTGTAACGTTGCCATTGCGCTAAGGTGTACTATGGTAAAAAAT | 50+6 |
| **WGAN15** | | AATTCTTTTGAGGTAAAGTGTTGCACACCCTGAAATGTTGTAGTATACTGGAGAGT | 50+6 |
| **WGAN16** | | AATTCTTATGATAAGTTCACTGTTGGCTAAAGGGAGTGTAAGTCATAATGAACTTT | 50+6 |
| **WGAN17** | | AATTCATGCACTGTTACCCTTTGTAATATTGTTCAGTATGGCCTATGCTACGCATT | 50+6 |
| **WGAN18** | | AATTCTTGATCAAAATTGAGTTTTATTATTTAAGCATATGGTATGTTTATAAACCT | 50+6 |
| **WGAN19** | | AATTCTGTTTGCATCTTGCGCTGAAACACTGCGCACCTGTGCTATACTATGCACAT | 50+6 |
| **WGAN20** | | AATTCAGGTAAGATACAATGTGAAGTAAAGATAAATCACGCCTATGCTAAAACAGT | 50+6 |
| **WGAN21** | | AATTCGTAAACTTTTGAAAATTAAATAAACTTTAAATTCTTGCTAAAATGGGCGTT | 50+6 |
| **WGAN22** | | AATTCTGTCAGGTTTTGAACTTGAAGAAGAAGTGGTGTGTGGCTATAATTGAGTTT | 50+6 |
| **WGAN23** | | AATTCTTTTGAAATCAAATTGTTATAACTAAAGTAAACTCTGCTATATTGCTCAAT | 50+6 |
| **WGAN24** | | AATTCTGTTTAAAAAAGCTGATTAAAATTTATTAATTTGTGCTATAATCCAGCATT | 50+6 |
| **WGAN25** | | AATTCACGTTGCGACAACATTTTGCGTATTCGCTAAAAGATGTTAGAATAATGCCT | 50+6 |
| **WGAN26** | | AATTCACGCACACTTTCACGTCGCAAAATGGTACACTATTGGCTATGCTACGCACT | 50+6 |
| **WGAN27** | | AATTCATTTACGTTGACCATTTGTAAAAATAAGCAATAATGGCTAAAATTAACCTT | 50+6 |
| **WGAN28** | | AATTCATTTCAAGATTAAGTTTTGATCCTGAACTAAAATGGGCTAAGTTACTGTCT | 50+6 |
| **WGAN29** | | AATTCGAGAACATGTTTGCGTTGAAAAACTTGTAAGGTTGGGCTAAAATAGCGTAT | 50+6 |
| **WGAN30** | | AATTCACGTTACAGAAATACGTGGTATTTGCGAAGTTTTGGGCTAGACTGGAGCGT | 50+6 |
| **WGAN31** | | AATTCTTACATTGTGAAACGTTTCACACTTGCGCATTATGTGCTATGCTGGGCAAT | 50+6 |
| **WGAN32** | | AATTCGTTTGCACTGAAAAGATTTGCGCTAGGGTGTAAGATGGTATGATACGTTTT | 50+6 |
| **WGAN33** | | AATTCTGGTAAAAAAAACTGTGGGAAAATTTGTTAAAACTTGGTAAAATAGCCTAT | 50+6 |
| **WGAN34** | | AATTCAAGTATCGCACACGGTGAACTTTGACTGAACTCCGCCTACGATGCGAATTT | 50+6 |
| **WGAN35** | | AATTCTATTGTTGTTGCGTCATAATGTTGTAAAGTTTTTAGGCTTATTCACGAATT | 50+6 |
| **WGAN36** | | AATTCTTTTGACGTGTTGACTTAAACGTGCAAGCAGGATTGCTAAACTGTACGCCT | 50+6 |
| **WGAN37** | | AATTCTTGTGCAGTTAATATTTGCAAAAATGGGTAAAAGCAGGTACAATAACTAAT | 50+6 |
| **WGAN38** | | AATTCATTGTGCTTTGCGCCTGACGTCTTAGCGGTATTGTACTATATTCCGCCGCT | 50+6 |
| **WGAN39** | | AATTCTTTGAAAGAAGACTATTGAGTTTGCAGGTTCTGAGGCTATACTTTGAATCT | 50+6 |
| **WGAN40** | | AATTCTTATGAAGAAAACGTTTTGAAAAAGGCATTAAAATTATACCATGAACATCT | 50+6 |
| **WGAN41** | | AATTCGCGAAGTCTGGAAGGATTAAATTCTTTGCTTTTGGACTACACCTCAAGACT | 50+6 |
| **WGAN42** | | AATTCCGGAAATTTGCCGAAAGCCACAACCTGTTAAGTTGCCTCTAATAGAAATCT | 50+6 |
| **WGAN43** | | AATTCCTTTTTGCGTAACAATTGATTTATTCGCATGGAAGTATAATACTCGTTTCT | 50+6 |
| **WGAN44** | | AATTCTTCTTAAGTAAAACGTTGAACTCTTCGATTTGCGATACTATCATTCTTCCT | 50+6 |
| **WGAN45** | | AATTCGAGAACTTCTATGGGTGGCTAATGTGACGAAAGAAGGTATGCTGGTCACCT | 50+6 |
| **WGAN46** | | AATTCATAAAAACCATGAAATTGCCACCTCGCCAATTTGATGTATGATTTTCCTGT | 50+6 |
| **WGAN47** | | AATTCTTGCGACGTTGAGGGTTTGCCATCTGCGCAATATATGGTTAAACATGATAT | 50+6 |
| **WGAN48** | | AATTCAAGAACTTGTGATCAAATAACTTTGTCGCAGAAACGTTTAAAATGAAGGCT | 50+6 |
| **WGAN49** | | AATTCCTGTAGATGTTTTCATGCAATTATGAATAATAATCTGATAAAGGATGACCT | 50+6 |
| **WGAN50** | | AATTCATGTTGCGTTGCGTTTTAATAAAGGCGTTTTCGTCGCCTATACTTCTTCCT | 50+6 |
| **WGAN51** | | AATTCTCTGCAAAATTAATGTTAATCGGTTCGTCTGAAAAAGCTATGTTAATAAAT | 50+6 |
| **WGAN52** | | AATTCCGGTTTTCTGCACGCTTTAAGCTAAGCAAAATCCGCCTATGATTAGCCTCT | 50+6 |
| **WGAN53** | | AATTCTTAAATGTAAGCATTGCGCTTCCTTACCAGAAAACTTTTAGAATAGTCCTT | 50+6 |

| | | | |
|---|---|---|---|
| **WGAN54** | | AATTCTTGAAACGTCTACCTTTGACTTTAATCTTTAGATTTTTCATCTTAACATAT | 50+6 |
| **WGAN55** | | AATTCCTAGGAGCGGTATTTCCTTACTTTCCTGTTATATGTTCTACTGTCTGATTT | 50+6 |
| **WGAN56** | | AATTCTTTAAGTTTGCGGGCATTAAATCACTTCTACGTCTGTTATGTTATGATAAT | 50+6 |
| **WGAN57** | | AATTCCGAATCACTAGATCTCTGGCATTGATTTAAATAAGATAAAAGTATGACTTT | 50+6 |
| **WGAN58** | | AATTCGAATTTACCTATCCTTTTATTTACCAAACTAAATTTCACAAGATATAATCT | 50+6 |
| **WGAN59** | | AATTCTTCTGCACCTGAAGGTTATACGTCAAGTGTTTATATTGTATCATGACACTT | 50+6 |
| **WGAN60** | | AATTCTCACTGAATAGAACATATTACGCGTTATCCTCACCAATATACTTTGTGGCT | 50+6 |
| **WGAN61** | | AATTCAACGAAATTTTGTATTTTTCAGAAGATACAGTCGGCAATATTTTGAAAACT | 50+6 |
| **WGAN62** | | AATTCTTAAGAACAATAATCACAGCTAAACATTGGCACTCATCTATATTAAGACTT | 50+6 |
| **WGAN63** | | AATTCTGAAAGATTTTTGCTGGGACGTCACCAAATAAATCGCTATATGATGTGTTT | 50+6 |
| **WGAN64** | | AATTCATTGGCGTCATTTTTTGACGTAGTCAAAACTCTGGCAGCATACTACGAATT | 50+6 |
| **WGAN65** | | AATTCACAGCTGCGTAGTTGTTGGCTTTTGCAAAACAAGGCCATATTCTTTAATAT | 50+6 |
| **WGAN66** | | AATTCGCTGATGAACAGTGTTGACGTTATTATGACGTTTCTCGTTAATCAATCCTT | 50+6 |
| **WGAN67** | | AATTCGCTTTCTCGACTAACGTATTGGTAATGCCAAATAAATTATTAAGTGAAAAT | 50+6 |
| **WGAN68** | | AATTCAGAACGCTTTCTTCTTTAACGCGTCATTAGGGCTTTTACGATGAAAGTGTT | 50+6 |
| **WGAN69** | | AATTCATATCGTAAAGATTTTACAAGTAAACGTTCGCGTTTGTTAAGAGAATTGTT | 50+6 |
| **WGAN70** | | AATTCTTTTCCGTGTCATTAATTGCTATTGGTTAAGTGGCATAACAATCTGCAACT | 50+6 |
| **WGAN71** | | AATTCTAAAAGTATCTGGATAATAACATCCTGATAAACCTGTATAAGTTGACCGCT | 50+6 |
| **WGAN72** | | AATTCTCTGCGACGAGTTCTTTCCTTTTTTCAAGCGGTTATGTCAAAATTCGCCAT | 50+6 |
| **WGAN73** | | AATTCAGGCGATACAACGATTTTATCAACGGCAGCATCTGTCATATACTATTAAAT | 50+6 |
| **WGAN74** | | AATTCTTAACAAAAAAAGCGTTCGCGGAATCATATTTATCACCGATACATCCACTT | 50+6 |
| **WGAN75** | | AATTCTGCAGTAACATCGCGTCGAAGCCTTTCTTCCTTTTTTGTACAATGGCTATT | 50+6 |
| **WGAN76** | | AATTCGTGCTATTGCTTGAAGTAGTATTGCTTTTTCACGCGAAAAAACCTTAGCCT | 50+6 |
| **WGAN77** | | AATTCATTATCCGGCTAAAATCTAGCAACTACACAAGACGATTTATAATGTATATT | 50+6 |
| **WGAN78** | | AATTCCGCCAGATGCGCCAATTTCTGTCAGTGGTCACGCGTGGCGCACTGACCTGT | 50+6 |
| **WGAN79** | | AATTCAAATCATCAACGTTTTTGCTTTTTGCGCGCCTCACCACTACGATAATTGAT | 50+6 |
| **WGAN80** | | AATTCGAAGTGTCGGCATGATACCAGCACGTTATTCTTCCTGTCAAAATCTTGCCT | 50+6 |
| **WGAN81** | | AATTCCTGCCCAGCATGCTCCCTGCCGATGTGAGCCGCCTGTTTATCCTGCGAAAT | 50+6 |
| **WGAN82** | | AATTCCCTTCTATCCGGGTTAAACCACCTTGATCCAGCGCTGCGAAGGTAAAGGCT | 50+6 |
| **WGAN83** | | AATTCGCGTTGGCGGCATTAGTGGCGCAGCAGAACGATTTGTCGATAATCCTACCT | 50+6 |

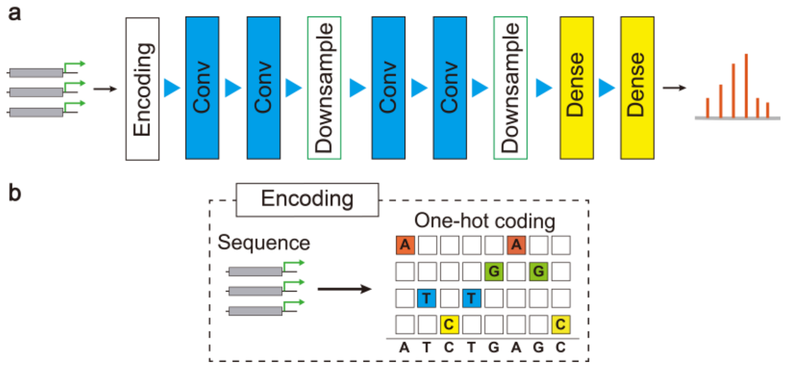


Figure S1. (a) The network structure of predictor network for predicting gene expression level. (b) Encoding process: DNA sequence is encoded into one-hot matrix.

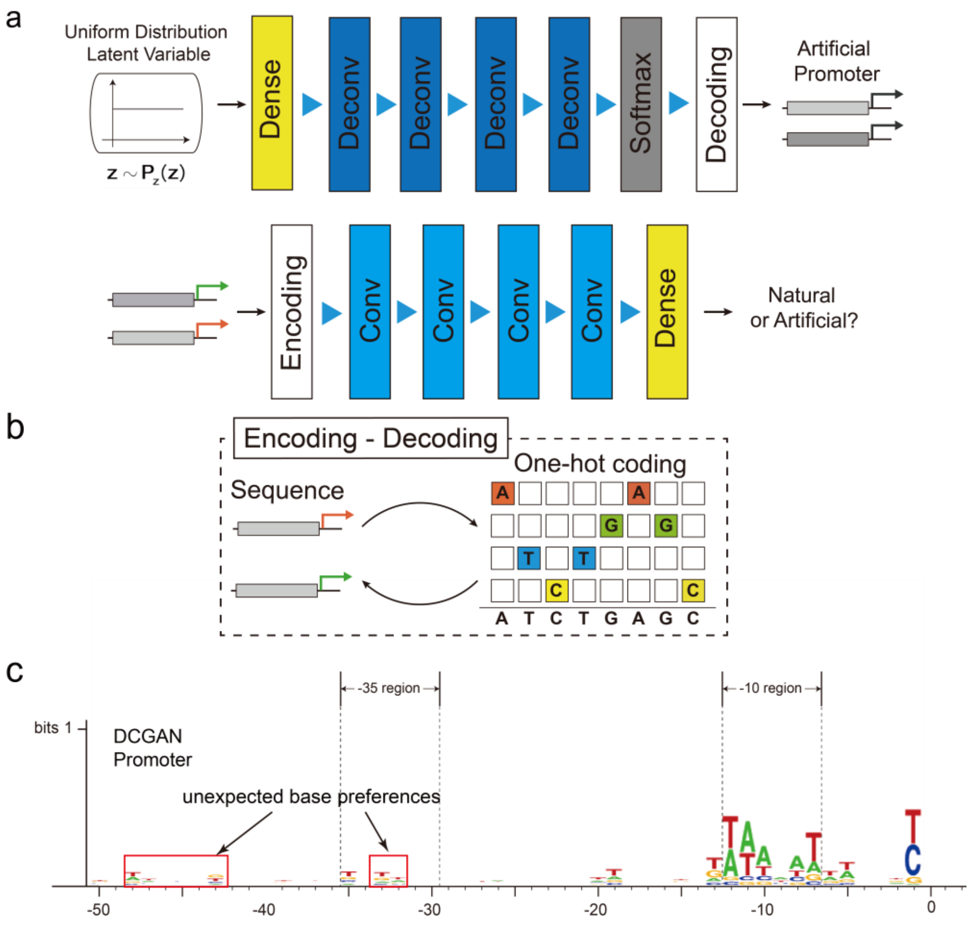


Figure S2. (a) The network structure of DCGAN. The input of generator is uniform distribution latent variable. (b) The encoding-decoding process: DNA sequence is encoded by one-hot matrix and the one-hot matrix is transformed back into DNA sequence by the maximum output of softmax function. (c) The DCGAN sequence logo of –1 ~ –50 region relative to transcriptional start site is shown in the figure. And the –10 region and –35 region are annotated.

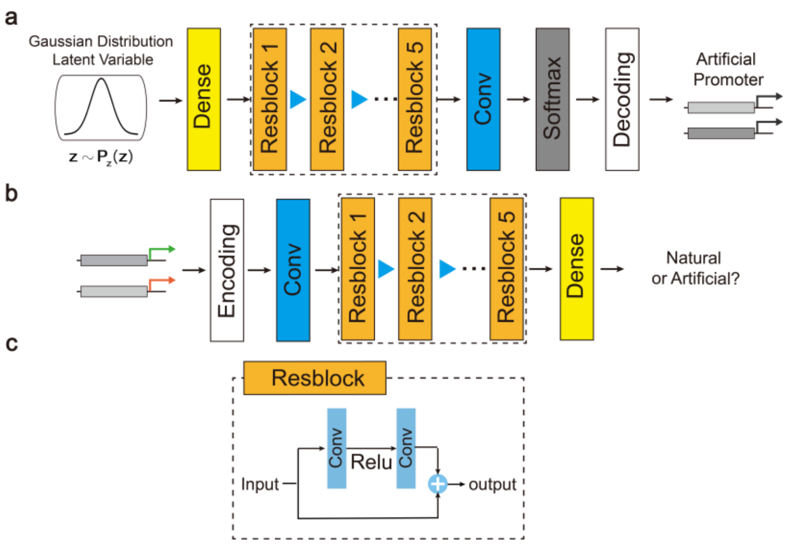


Figure S3. The structure of WGAN-GP language model. (a) Generator network structure. The input is gaussian distribution latent variable. (b) Discriminator network structure. (c) The resblock structure.

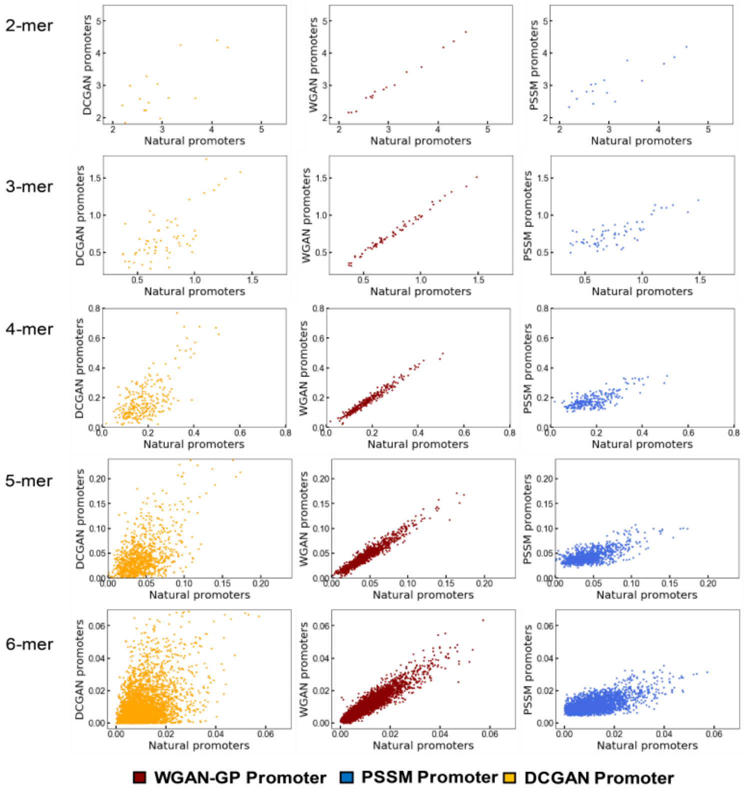


Fig S4. The scatter plot of k-mer frequency between model-generated promoters and natural promoters from k = 2 to k = 6. The model-generated promoters contain DCGAN (orange), WGAN-GP (red), PSSM promoters (blue).

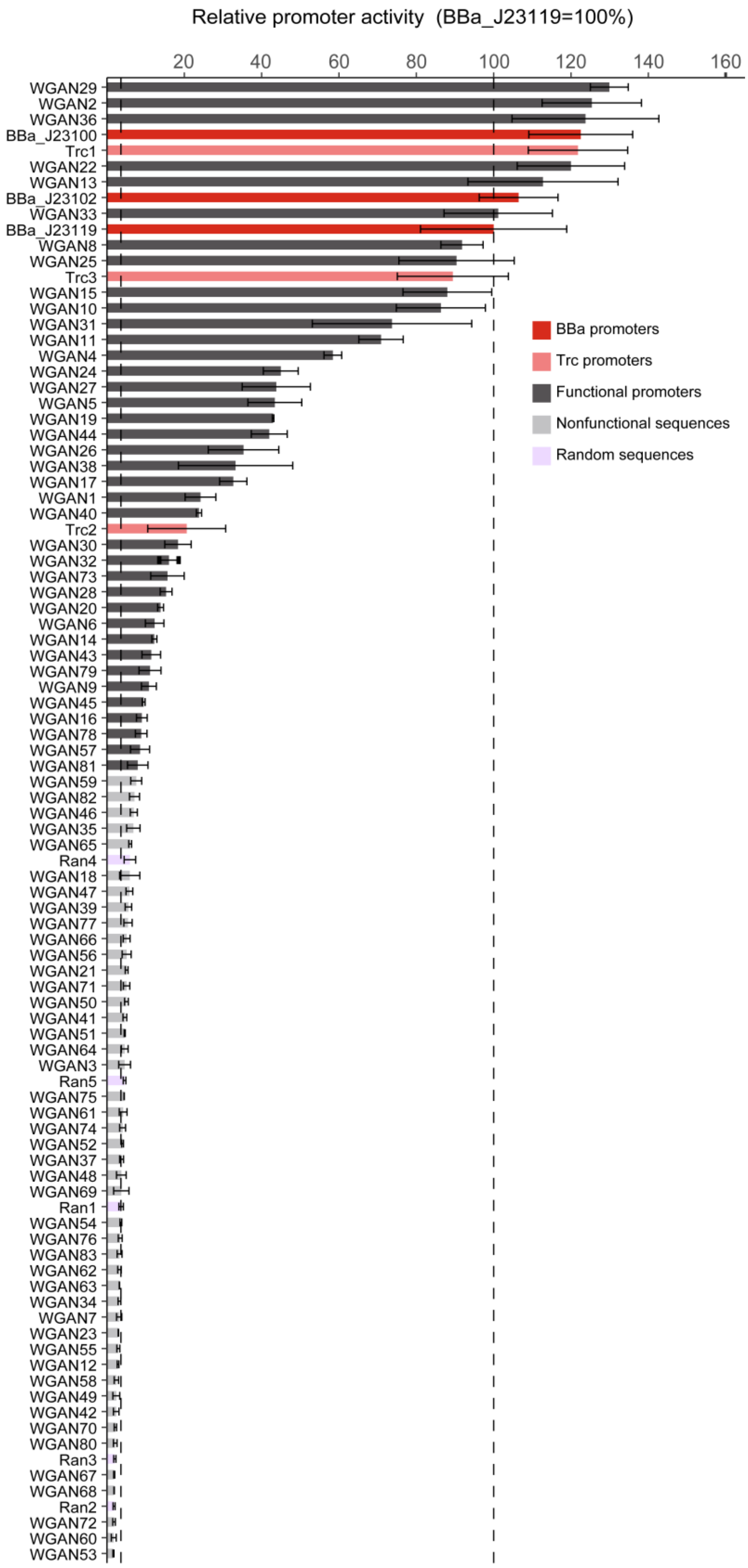


Fig S5. The relative activity of 83 synthetic promoters designed in silico. BBa J23119 (dark red) and Trc (light red) are wild-type promoters. BBa_J23100, BBa_J23102 and Trc2, Trc3 are their highly expressed mutants. Functional promoters (grey) and non-functional promoters (light grey) and 5 random sequences (purple bars) are shown. The dashed line represents the activity of BBa_J23119 (right) and average activity of five negative control promoters(left) respectively.

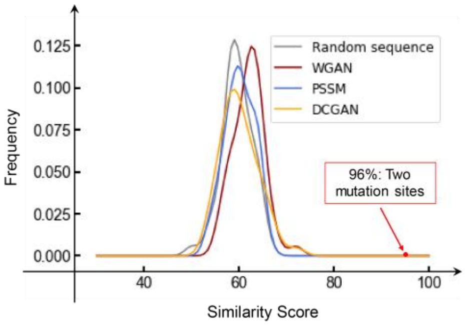


Fig S6. The similarity distribution of random sequence (grey), WGAN promoters (red), PSSM promoters (blue) and DCGAN promoters (orange). Different from methods based on mutagenesis, which only mutant one or two sites in promoter sequences, WGAN promoters show low sequence similarity with natural promoters.

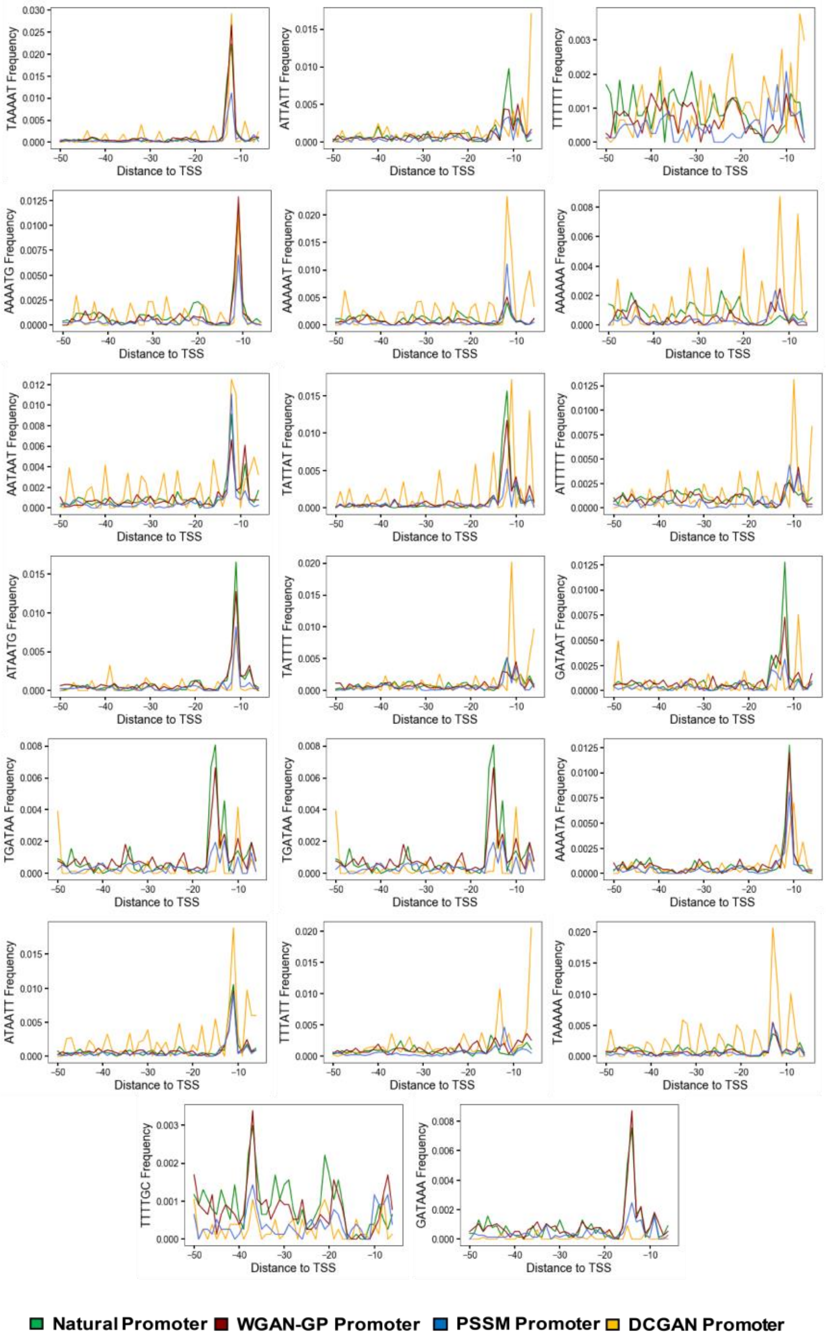


Figure S7. The position distribution relative to TSS of top 20 most frequently occurring 6-mers are shown in the figure. These 6-mers in natural (green), WGAN-GP (red) PSSM (blue) and DCGAN (orange) promoters show different location preference patterns.
